# Prenatal Dibutyl Phthalate Exposure Dysregulates Fetal-Placental Vascular Function and Placental Vasculature-Specific Lipid Metabolism

**DOI:** 10.64898/2026.07.11.732137

**Authors:** Xinran Yang, Dilen Kadlec, Jordan Schultz, Zelieann R. Craig, Chi Zhou

## Abstract

**Introduction:** In humans, prenatal dibutyl phthalate (DBP) exposure is associated with increased risks of adverse fetal outcomes as well as metabolic and cardiovascular diseases in the offspring in a fetal sex-specific manner. However, mechanisms underlying these prenatal DBP exposure-associated adverse fetal/offspring outcomes are unclear. We **hypothesize** that environmentally relevant low-dose prenatal DBP exposure dysregulates fetal-placental vascular function and lipid metabolism in a fetal sex-specific manner, thereby impairing placental efficiency and programming adverse offspring metabolic outcomes.

**Methods:** Female CD-1 adult mice (8-10 weeks) were orally dosed with vehicle or an environmentally relevant low-dose DBP (0.1μg/kg/day) daily from 30 days pre-pregnancy through gestational day (GD) 18.5. Fetal-placental vascular hemodynamics of these dams were examined using high-frequency ultrasound at multiple timepoints. The effect of prenatal environmentally relevant low-dose DBP exposure on placental efficiency, spatial transcriptomic profiles, lipid homeostasis, and placental vascular endothelial cells function in male and female fetuses were evaluated at gestational day (GD) 18.5.

**Results:** The prenatal low-dose DBP exposure dysregulated the fetal-placental vascular hemodynamic indices from mid-to late gestation. DBP exposure impairs placental efficiency in male, but not female placenta at GD18.5. Further, female placentas exhibited fetal labyrinth vasculature-specific transcriptomic adaptations that preserves placental efficiency and endothelial function. In contrast, male placentas exhibited minimum transcriptomic adaptation, together with compromised placental efficiency and endothelial function associated with lipotoxic lipid profile.

**Conclusions:** In conclusion, prenatal low-dose DBP exposure dysregulates placental vascular function and lipid homeostasis in a fetal sex-specific manner, with male fetuses being more susceptible to DBP exposure.

## Introduction

Phthalates are known to adversely impact cardiovascular function in humans^1–4^. Despite the gradual decrease in legacy phthalate use in recent years, dibutyl phthalate (DBP), a phthalate congener, is still detected in human biofluids^3, 5–9^. Human DBP exposure are estimated around 0.1-233µg/kg/day through consumer products, occupational exposure, and medication coatings^10–12^. In general population, women show greater burden of DBP than men (indicated by elevated urinary concentration of mono-n-butyl phthalate [MBP], the primary metabolite of DBP and marker of DBP exposure)^3, 7–9^. DBP exposure was also reported in most mothers and fetuses during pregnancy examined across multiple biomonitoring studies^13–15^. In humans, DBP exposure during pregnancy is associated with pregnancy complications as well as metabolic and cardiovascular diseases in the offspring^13–21^. To date, mechanisms underlying prenatal DBP exposure-associated adverse pregnancy/fetal outcomes are unclear.

DBP and its main metabolite MBP are low molecular weight phthalate congeners that can regulate lipid metabolism and can cross the maternal-fetal interface^22–24^. In humans, prenatal phthalate exposure is reported to be associated with increased risks of intrauterine growth restriction, with male fetuses more susceptible than female fetuses^16^. Prenatal DBP exposure is associated with risks of large for gestational age at birth and higher body mass index and waist circumference during childhood in a fetal-sex specific manner, with stronger association in male offspring^25, 26^. Our previous report in mice has demonstrated that environmentally relevant prenatal DBP exposure fetal sex-specifically dysregulates the late gestational placental lipid deposition and placental efficiency in a fetal sex-specific manner^27^, indicating potential role of lipid metabolism in DBP-induced adverse offspring outcomes. However, the mechanism linking prenatal DBP exposure to altered placental lipid deposition and potential metabolic dysfunction in the offspring remain not fully understood.

To date, most existing DBP studies investigating environmentally relevant dosages have been focused on gonadal toxicity in non-pregnant mice, while studies focused on pregnancy/fetal outcome studies typically using dosages substantially higher than human environmental exposure levels^18, 28–32^. Although environmentally relevant DBP exposure is reported to disrupt various gonadal processes (e.g., folliculogenesis, steroidogenesis, cell cycle, and apoptosis)^29, 33–37^, the effects of environmentally relevant low-dose DBP exposure on placental and fetal vascular development and function remain largely unexplored. In addition, despite epidemiological evidence linking prenatal phthalate exposure to adverse fetal growth outcomes in a sexually dimorphic manner in human, the mechanistic effects of environmentally relevant prenatal DBP exposure on placental-fetal vascular and metabolic function across fetal sex remain to be elucidated.

Previous human studies have linked prenatal DBP exposure (indicated by urinary MBP levels) with altered transcriptomic profiles in bulk placental tissue^38–42^. However, these studies evaluated multiple co-occurring phthalates making it difficult to establish a clear causal relationship between DBP exposure and specific placental transcriptomic changes. Further, as placental tissue consists of multiple cell types, the heterogeneity of placental tissues also contributes to the highly varied bulk placenta transcriptomic profiling data in the literature. As the morphological structure constructed by these different cell types is key to placental vascular/endothelial function and fetal development, transcriptomic profiling analyses that capable of mapping specific gene expression changes to morphological structures are essential.

In this study, we tested the hypothesis that environmentally relevant low-dose prenatal DBP exposure dysregulates fetal-placental vascular function and lipid metabolism in a fetal sex-specific manner, thereby impairing placental efficiency and programming adverse offspring metabolic outcomes. Specifically, we examined the effect of environmentally relevant prenatal low-dose DBP exposure on fetal-placental vascular hemodynamics. We further determined the effect of prenatal environmentally relevant low-dose DBP exposure on placental efficiency, spatial transcriptomic profiles, lipid homeostasis, and placental vascular endothelial cell function in male and female fetuses at gestational day (GD) 18.5.

## Methods

### Data Availability Statement

The authors declare that all supporting data are available within the article and it’s corresponding publicly accessible datasets (NCBI Gene Expression Omnibus [GEO] Data Series # GSE333401).

### Ethical Approval

All animal experiments followed guidance of the Animal Research: Reporting of In Vivo Experiments (ARRIVE) guidelines^43^, the Guide for the Care and Use of Laboratory Animals by National Research Council^44^, and were performed with the approval of the University of Arizona Institutional Animal Care and Use Committee.

### Animals

Adult (8-10 weeks, weight 32±2 gram) female CD-1 mice were purchased from Charles River Laboratories (Charles River, California) and housed in single-use, BPA-free cages at the University of Arizona Animal Facility with ad libitum water and food (14 L:10 D light cycle, 22±1°C temperature). After at least 1 week of acclimation period, mice were randomly assigned to receive daily oral treatment of vehicle (tocopherol-stripped corn oil; MP Biomedicals, OH; vehicle; control group) or DBP (tocopherol-stripped corn oil supplemented with 0.1μg/kg/day of DBP; Sigma-Aldrich, MO; referred as DBP) as previously described^27^. Based on reports of human exposure estimates (range 0.1-233μg/kg/day) and previous studies (0.1-100µg/kg/day), this environmentally relevant low DBP dosage does not cause systemic toxicity nor weight gain as evidenced by daily normal behavior, body condition, and weight gain pattern in both non-pregnant and pregnant CD-1 mice^27, 29, 30, 45, 46^.

### Daily Oral DBP Dosing in Non-pregnant and Pregnant Mice

Adult female CD-1 mice were mated with naïve male CD-1 mice without DBP exposure after 30 days of daily oral treatment of vehicle (control [CT] group; n=18) or DBP (n=18) as previously described^27^. The same daily oral treatment of CT and DBP groups continued through pregnancy until gestational day (GD) 18.5. Each mouse was weighed daily to monitor weight-gain pattern throughout treatment period. Mice were euthanized at GD18.5 under pre-sedation with inhaled isoflurane overdose followed by decapitation. Weight of individual fetuses and matching placentas were measured for all fetuses in each litter. Placental tissues were collected and immediately processed for downstream spatial transcriptomics, lipidomics, microvasculature endothelial cell isolation, and total RNA isolation (**Fig.1**).

**Figure. 1.**
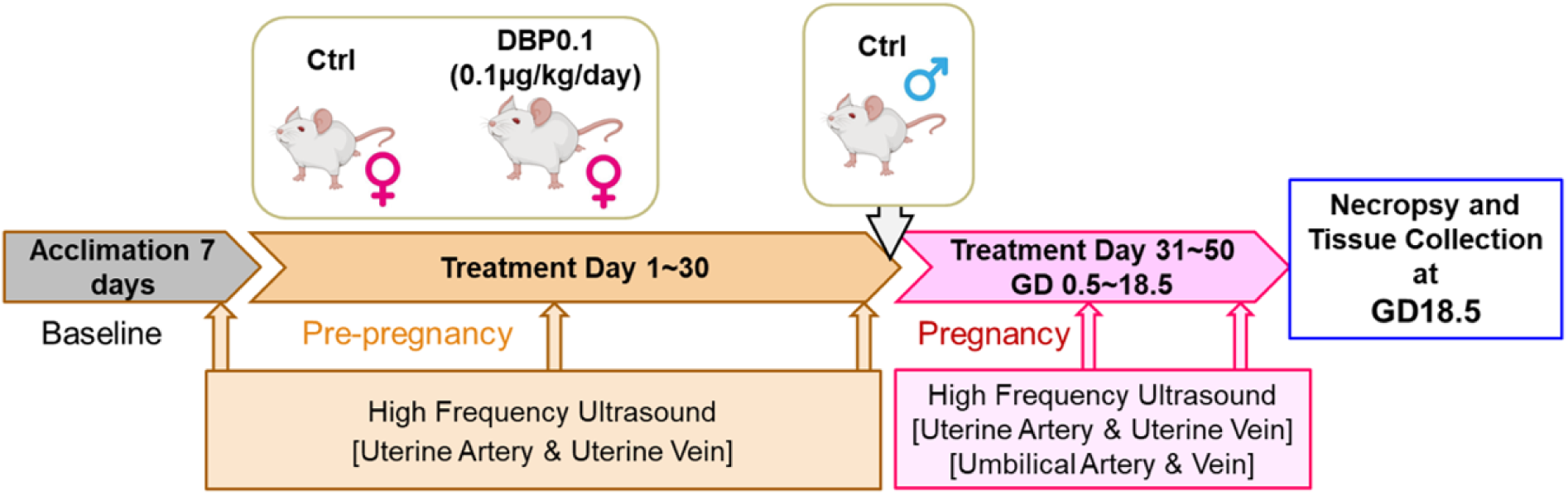
Experimental design for animal dosing and physiological measurements.

### High-Frequency Ultrasound Imaging Assessment of Maternal Uterine-Placental and Fetal-Placental Hemodynamics

Hemodynamics of maternal uterine arteries, maternal uterine veins, and fetal umbilical arteries were assessed in mice using high-frequency Doppler ultrasound using a Vevo 3100 imaging system (FUJIFILM Visual Sonics, Inc; CT: n=10, DBP: n=12). Maternal uterine artery and uterine vein Doppler wave-form measurements were obtained at baseline/day 0, day 15, and day 30 of the pre-pregnancy exposure period, as well as GD11.5, and GD18.5 during pregnancy. Fetal umbilical artery Doppler wave-form measurements were obtained from 2 fetuses per dam at GD11.5 and GD18.5. The high-frequency ultrasound Doppler wave-form measurement was performed as previously described^47^. In brief, mice were anesthetized and placed in the supine position on a temperature-controlled (37-38°C) imaging platform. Abdominal hair was removed, and prewarmed ultrasound gel was applied as the acoustic coupling medium. Maternal body temperature and heart rate were maintained during image acquisition. Target vessels were identified using B-mode and color Doppler imaging. Peak systolic velocity (PSV), end-diastolic velocity (EDV), and resistance index (RI; calculate as RI=(PSV-EDV)/PSV) were assessed in each vessel using the VevoLab software (FUJIFILM Visual Sonics, Inc).

### Fetal, Placental, And Placental Efficiency Distributions at GD18.5

Individual fetal weight and matching placental weight of all conceptuses from each dam at GD18.5 was collected (CT: n=14 dams, DBP: n=14 dams). The distributions of fetal weight, placental weight, and placental efficiency (calculated as fetal weight/placental weight ratio) of conceptuses were classified into low (0^th^-10^th^ percentile), intermediate (10^th^-90^th^ percentile), and high (90^th^-100^th^ percentile) categories based on all control conceptuses. These thresholds were adapted from the conventional clinical classification of human neonatal weight as small, normal, or large for gestational age, respectively^48, 49^.

### Spatial Transcriptomic Profiling of GD18.5 Placentas

Spatial transcriptomic sequencing was performed with female (F) and male (M) mouse placentas from control (control female [CT-F]: n=3, control male [CT-M]: n=4) and DBP exposed (DBP-F: n=3, DBP-M: n=3) dams using the Visium Spatial Gene Expression Slide and Reagent Kit (10x Genomics) followed by high-throughput next-generation sequencing as described^50^. In brief, GD18.5 placental tissues were embedded in OCT compound in an isopentane/liquid nitrogen (LN_2_) bath immediately after necropsy. Fresh frozen placental tissue sections (10μm thick) were fixed on the Visium HD Spatial Gene Expression Slide and bind with corresponding spatial bar-coded units followed by H&E staining following the manufacturer’s instructions. After the H&E staining, Visium on-slide library construction was performed with spatial bar-coded units binding tissue sections followed by Illumina NovaSeq high-throughput transcriptomic sequencing analysis as described^50^. To identify morphological structure-linked transcriptomics profile changes, spatial transcriptomic data were analyzed using the Space Ranger software (10X genomic spatial transcriptomics data analysis platform), RCTD software^51^ (cell type annotation), and the Suerat^52^ package. The cell type composition of each spatial bar-coded unit was annotated using the RCTD^51^ software. The transcriptomic data of each spatial bar-coded unit was mapped to the corresponding Visium H&E staining image using the Suerat^52^ package. The spatial bar-coded units-linked transcriptomic profiling data were integrated using the Harmony^53^ package, clustered based on their transcriptomic profile using Seurat^52^ package, and proceed with the Uniform Manifold Approximation and Projection (UMAP)^54^ analysis for two-dimensional visualization of resulting clusters in R. The morphological structure represented by each cluster was determined based on RCTD-annotated cell-type composition and the spatial distribution of spatial bar-coded units in the tissue section. Pseudo-bulk transcriptomic and functional genomics analysis was further performed using Suerat^52^, edgeR^55^, and Ingenuity Pathway Analysis (IPA) software to identify differentially expressed (DE) genes and pathways at the biological sample/treatment group level^50, 56–58^.

### Lipidomic Profiling of Mouse Placenta

Ultra-high performance liquid chromatography-tandem mass spectrometry (UHPLC-MS) analysis of 12 classes of lipids (total of 136 molecules) that are critical to lipid metabolism and transport was performed in individual placental tissues following the established workflow^59^ by the University of Arizona Analytical Chemistry Shared Resource (ACSR). In brief, individual GD18.5 placenta tissue was snap frozen in LN_2_ immediately after necropsy and stored in -80°C. The snap frozen tissue was then homogenized in liquid nitrogen (LN_2_), followed by an adapted isopropyl alcohol: ethyl acetate (1ml) extraction^59^ and 10,000 x g centrifugation for 10min at 4°C. Supernatant was then dried under nitrogen and resuspended in 100μl of 45:40:5 Isopropyl alcohol: Acetonitrile: Water (5 mM ammonium formate,0.1% Formic acid). 1μL of extract was injected into the Thermo Vanquish Duo UHPLC with the C18+ (150x2.1mm, 1.5μm) separate system. Mass spectrometry (MS) detection was performed using Thermo Exploris 480. Lipid annotations were assigned based on MS signals compared against metabolome databases (MZcloud, Lipidmaps, HMDB, Metlin) using Lipid Search5 (Thermo Scientific™).

### Isolation And Culture of Primary GD18.5 Placental Vasculature Endothelial Cells (PVECs)

After necropsy, individual placental tissues were immediately minced into small pieces (<0.5mm) and digested in Hanks’ balanced salt solution supplemented with 0.25% collagenase B (Roche) for 1 hour at 37°C. The digested tissue suspension was then centrifuged at 1,200 × g at 4°C. The cell pellet was resuspended in endothelial complete growth cell culture medium (ECM, ScienCell #1001) and seeded into 35mm cell culture dishes. Cells were incubated at 37°C in a humidified incubator with 5% CO_2_ for 4-6 hours followed by 3 washes with ECM to remove tissue debris and non-adherent cells. The purity of isolated PVECs were assessed using the AcLDL uptake assay as we previously described^56–58^. In brief, cells were incubated with ECM supplemented with 5μg/mL of Alexa Fluor™ 488 Conjugated Acetylated Low-Density Lipoprotein (Alexa Fluor™ 488 AcLDL; Invitrogen™ Cat# L23380) for > 6 hr at 37°C, followed by 10min of 10ug/mL Hoechst 33342 (Thermo Scientific™, Cat#62249; staining the nuclei of cells) in ECM. The stained live cells were imaged using the Cytation5 multi-mode imager (Agilent) using the Gen5 software. Total cells and cells exhibiting uptake of AcLDL were counted. Cells incubated without Ac-LDL were served as negative controls. The purified PVECs were cultured for up to 14 days with daily medium changes. Cell growth and confluency were assessed under microscope every 2 days.

### RNA Isolation and RT-qPCR Analysis of Anti-apoptotic, Senescence, and Telomere-Maintenance Related Genes in GD18.5 Placenta

After necropsy, GD18.5 placental tissue samples were snap frozen in liquid nitrogen, stored at -80°C, and then ground to powder in liquid nitrogen followed by RNA isolation. Total RNA samples were isolated from GD18.5 placentas using the RNeasy Mini Kit (Qiagen, Valencia, CA). The concentration and quality of each RNA sample were assessed using a NanoDrop™ND-1000 spectrophotometer (NanoDrop Technologies, Wilmington, DE) and Agilent 2100-bioanalyzer (Agilent Technologies, Santa Clara, CA)^56–58^. Only RNA samples with a high purity (1.9 < OD260/280 < 2.1) and high RNA integrity number (RIN > 8) were utilized in this study.

RT-qPCR analysis of *Bcl2* (anti-apoptosis marker^60^), *Cdkn1a* (also known as P21, senescence/cell-cycle arrest marker^61^), and telomere maintenance-associated markers (*Dkc1*^62^ and *Tert*^63^) was performed in female and male placental tissues from CT and DBP groups (CT-F: n=9, CT-M: n=10, DBP-F: n=8, DBP-M: n=8) as previously described^56–58^. In brief, small RNA fragment enriched total RNA isolated from each sample was reverse transcribed into cDNA using a miScript II RT Kit (Qiagen). RT-qPCR was performed using QuantiTect SYBR Green PCR Kit (Qiagen) and commercially available Primer Assays (**Table.1**) using a StepOne^Plus^ qPCR system (Life Technologies, Carlsbad, CA). Efficiencies of all target and control assays were between 90% and 110%. Data were normalized to endogenous control genes (*Ubc* and *Polr2a*). The normalized data were then analyzed using the 2^-ΔΔCT^ method^58, 64^.

**Table 1.** List of RT-qPCR Assays.

| **Gene Symbol** | **NCBI Entrez Gene ID** | **Official Full Name** | **RT-qPCR Primer Assay type** | **Vendor** | **Catalog Number** | **Qiagen GeneGlobe ID** |
| --- | --- | --- | --- | --- | --- | --- |
| Bcl2 | 12043 | B Cell Leukemia/Lymphoma 2 | Gene of Interest | Qiagen | 249900 | QT02392292 |
| Cdkn1a | 12575 | Cyclin Dependent Kinase Inhibitor 1a | Gene of Interest | Qiagen | 249900 | QT00137053 |
| Dkc1 | 245474 | Dyskeratosis Congenita 1, Dyskerin | Gene of Interest | Qiagen | 249900 | QT01199387 |
| Tert | 21752 | Telomerase Reverse Transcriptase | Gene of Interest | Qiagen | 249900 | QT00104405 |
| Polr2a | 20020 | Polymerase (RNA) II (DNA Directed) Polypeptide A | Housekeeping Gene | Qiagen | 249900 | QT00197757 |
| Ubc | 22190 | Ubiquitin C | Housekeeping Gene | Qiagen | 249900 | QT00245189 |

### Statistical Analyses

SigmaPlot (Systat Software., San Jose, CA) and GraphPad Prism (GraphPad Software, Boston, MA) softwares were used for statistical analyses. Comparisons among groups across multiple time points were performed with two-way repeated measures ANOVA with post hoc Benjamini and Hochberg False Discovery Rate (FDR)^56, 58^ adjustments to correct multiple testing (FDR adjusted *P<0.05* considered significant). Two group comparisons at a single time point were performed with t-test (*P<0.05* considered significant). Comparison of categorical outcomes between CT and DBP groups was performed with Chi-square test or Fisher’s exact test, as appropriate (*P<0.05* considered significant). Comparisons among females and males from the CT and DBP groups at a single time point were performed with two-way ANOVA with FDR post hoc p-values adjustments for multiple testing (FDR adjusted *P<0.05* considered significant). Spatial transcriptomics and functional genomics data were analyzed using edgeR with FDR^56, 58^ adjustments to correct for multiple testing (FDR adjusted *P<0.05* considered significant).

## Results

### Effect of Environmentally Relevant DBP Exposure on Maternal Uterine Vascular Adaptation During Pregnancy

Our high-frequency ultrasound analysis data have shown that environmentally relevant prenatal DBP exposure does not affect the heart rate, and Doppler wave form patterns of maternal uterine arteries and veins in non-pregnant mice throughout the 30 days of treatment (**Fig.2**). We observed uterine vascular hemodynamic changes that consistent with the expected vascular adaptation from mid (GD11.5) to late gestation (GD18.5) in CT mice (**Fig.2A-D**). Specifically, the heart rate, peak systolic velocity (PSV), and end-diastolic velocity (EDV) of uterine veins and arteries maintained at levels similar to pre-pregnancy period at GD11.5 but significantly increased by GD18.5. In addition, the resistance index (RI) of uterine veins and arteries maintained at levels similar to pre-pregnancy period at GD11.5 but significantly decreased by GD18.5. These data are consistent with the expected increase in uterine-placental blood flow from mid- to late-gestation to support the rapid fetal growth by late-gestation in mice.

**Figure. 2.**
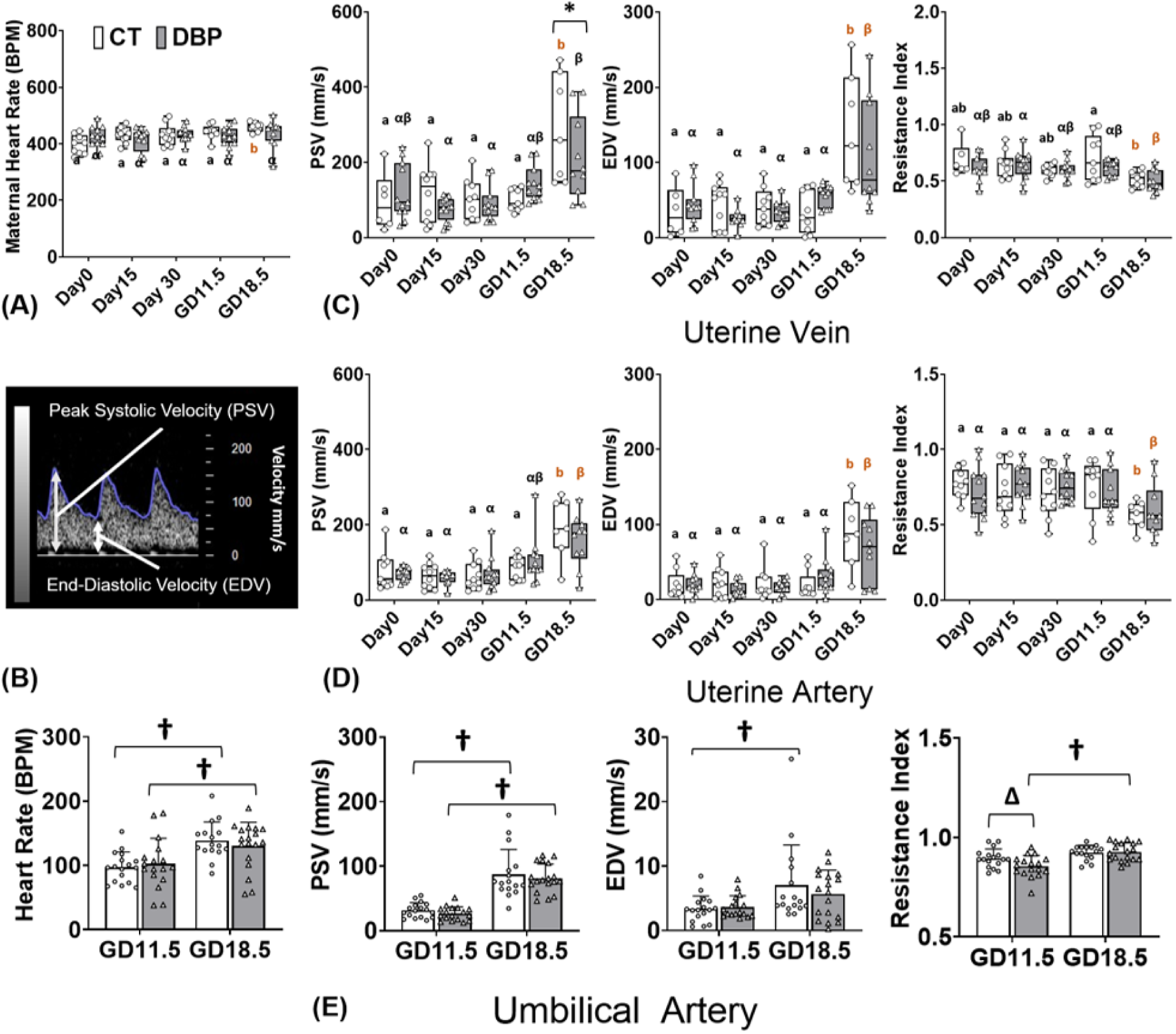
Effect of Low-Dose DBP Exposure on Maternal Uterine Vascular Adaptation (A-D) and Fetal-Placental Vascular Adaptation (E) During Pregnancy. (A) Maternal heart rate. (B) Representative hemodynamic measurement. PSV, EDV, and RI in maternal uterine vein (C) and artery (D). (E) PSV, EDV, and RI in fetal umbilical artery. (A-D): Different small letters (a vs. b in CT, α vs. β in DBP) represent significant difference across different timepoint within the same treatment group (CT or DBP); * Differ between CT and DBP groups within the same time point (P<0.05). Each data point represents the data from an individual dam. (E): **^Δ^** Differ between DBP and CT within the same time point; **^†^** Differ between GD11.5 and GD18.5 within the same treatment group. Each data point represents the data from an individual fetus. (CT: n=10 dams, DBP: n=12 dams).

Environmentally relevant prenatal DBP exposure at 0.1μg/kg/day did not significantly affect the hemodynamic parameter during the 30 days of pre-pregnancy treatment. However, the DBP exposure attenuated the heart rate increase at GD18.5. In addition, the PSV of maternal uterine veins elevated slightly at GD11.5 compared to CT dams while it later reached a significantly lower PSV at GD18.5 than their CT counterparts.

### Environmentally Relevant Prenatal DBP Exposure Dysregulate Fetal Umbilical Vascular Adaptation During Pregnancy

Our high-frequency ultrasound analysis of fetal umbilical arteries observed hemodynamic changes that were consistent with the expected vascular development from mid (GD11.5) to late gestation (GD18.5) in fetuses from CT dams (**Fig.2E**). Specifically, the fetal heart rate, umbilical artery PSV, and umbilical artery EDV all significantly increased from GD11.5 to GD18.5, while the RI maintained similar between GD11.5 to GD18.5 in fetuses from CT dams. The prenatal DBP exposure does not significantly affect the fetal heart rate and the umbilical artery PSV. However, the prenatal DBP exposure attenuated the EDV increase from GD11.5 to GD18.5 and lead to relatively lower RI at GD11.5 as well as increased RI from GD11.5 to GD18.5.

### Environmentally Relevant Prenatal DBP Exposure Causes Fetal Sex-Specific Shifts in Placental Efficiency Distributions at GD18.5

Compared to the CT animals, prenatal DBP exposure significantly shifted the fetal weight, placental weight, and placental efficiency distribution among fetuses in a fetal sex-specific manner (**Fig.3**). Specifically, female fetuses with prenatal DBP exposure were significantly overrepresented in the intermediate (10^th^-90^th^ percentile) category in fetal weight (91% vs. 78%, P=0.023), placental weight (94% vs. 79%, P=0.009), and placental efficiency (calculated as fetal weight to placental weight ratio; 92% vs 79%, P=0.019) compared with CT group. The proportion of female fetuses with high (90^th^-100^th^ percentile range) and low (0^th^-10^th^ percentile range) fetal weight, placental weight, and placental efficiency are decreased in DBP group but did not reach statistical significance. Prenatal DBP exposure did not significantly alter the distribution of fetal weight or placental weight of male fetuses. However, male fetuses with prenatal DBP exposure were overrepresented (25.5% vs. 10%; P=0.008) in the low placental efficiency category (0^th^-10^th^ percentile) and underrepresented (1% vs. 10%; P=0.007) in the high placental efficiency category (90^th^-100^th^ percentile) compared with counterparts from controls.

**Figure. 3.**
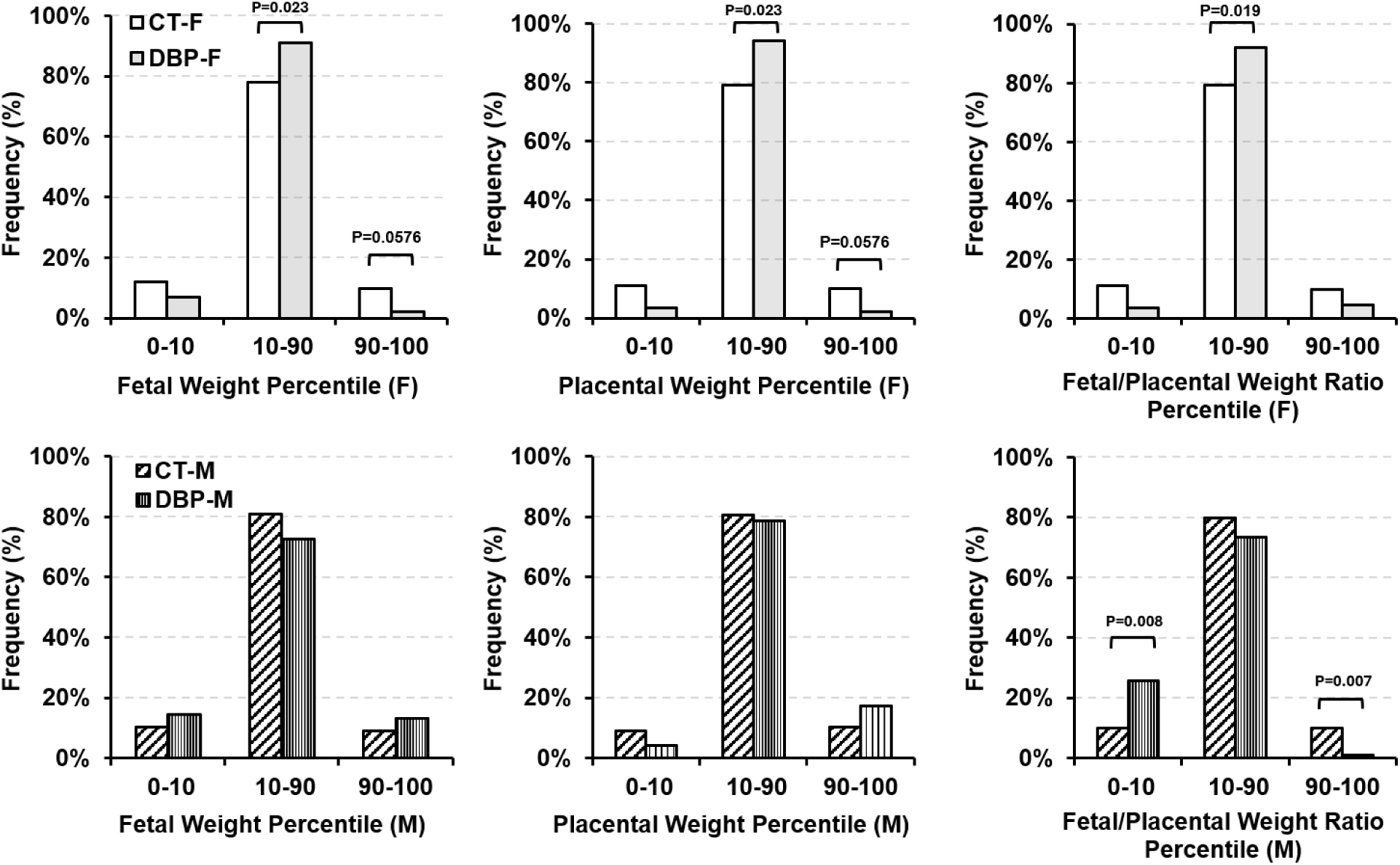
Effect of Low-Dose DBP Exposure on Distribution of Fetal Weight, Placental Weight, and Fetal/Placental Weight Ratio Among Male and Female Fetuses at GD18.5. (CT-F: n=91 fetuses (14 dams), DBP-F: n=89 fetuses (14 dams), CT-M: n=89 fetuses (14 dams), DBP-M: n=98 fetuses (14 dams).

### Prenatal DBP Exposure Induces Fetal-Sex Specific Transcriptomic Profile Dysregulation in Specific Fetal Labyrinth Vascular Regions

Our spatial transcriptomic profiling analysis revealed that environmentally relevant prenatal DBP exposure dysregulates the placental transcriptomic profile in a region- and fetal sex-specific manner (**Fig.4A-F**). We obtained a total of 772,584,586 reads from 22,246 spatial bar-coded units of the 13 placentas (CT-F: n=3, CT-M: n=4, DBP-F: n=3, DBP-M: n=3), with an average of 6,652 genes detected in each spatial bar-coded unit. Spatial transcriptomic bioinformatics analysis further revealed 33 clusters of spatial bar-coded units based on their transcriptomic profiles (**Fig.4A-C**). RTCD cell type deconvolution analysis revealed 16 cell populations observed among all placentas (**Fig.4D**). The RTCD cell type compositions of these 33 clusters are in agreement with the physical location of these clusters relative to the major placental morphological compartments (decidua: cluster 5, 12, 14, 16, 23, 25, 30; junctional zone: cluster 9, 10, 11, 15, 18, 22, 26, 28; labyrinth: cluster 0, 1, 2, 3, 4, 6, 7, 8, 13, 17, 19, 20, 21, 24, 27, 29, 31, 32).

**Figure. 4.**
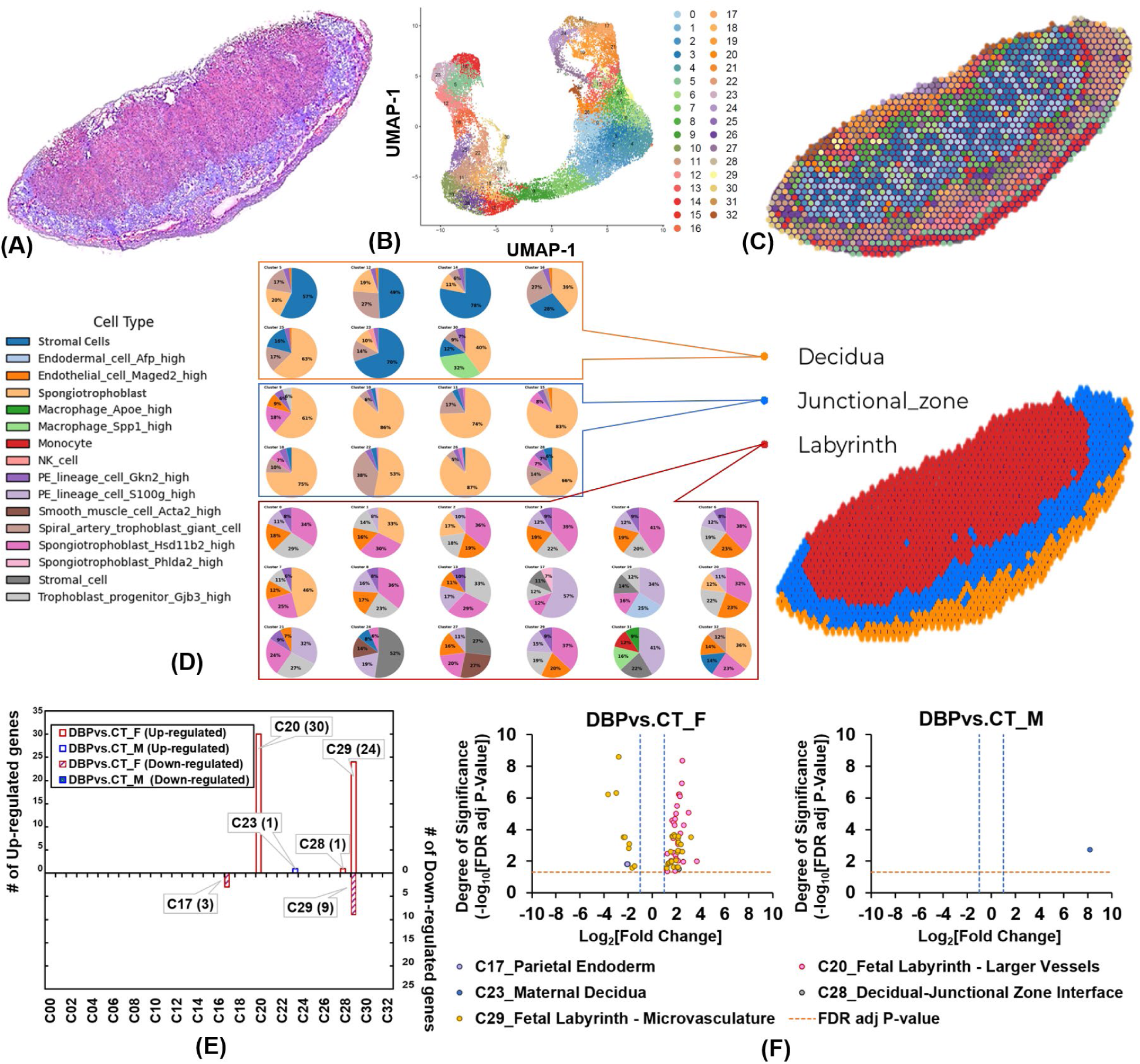
Effect of Low-Dose DBP Exposure on Female and Male Placental Spatial Transcriptomic Profiles at GD18.5. (A) H&E staining showing the morphological structure of the placenta tissue. (B) Clustering of spatial transcriptomic data based on gene expression profile. Method: UMAP-assisted harmony. Each dot represents the transcriptomic profile of one bar-coded unit on tissue sections. (C) Spatially resolved clusters that were mapped with the H&E-staining image of the placenta tissue section. Each dot corresponds to one bar-coded unit of clusters shown in (B). (D) RTCD cell type annotation and their morphological structure annotation. (E) Bar chart showing number DBP dysregulated genes in female and male placentas at the biological samples/groups level among clusters in pseudo-bulk analysis. (F) Volcano plot showing DBP dysregulated genes in female and male placentas at the biological samples/groups level among clusters in pseudo-bulk analysis. (n=3-4 per fetal sex/group).

Further pseudo-bulk gene expression analysis revealed that the environmentally relevant prenatal DBP exposure only dysregulated the transcriptomic profiles of 68 genes among 5 clusters (cluster 17, 20, 23, 28, 29) out of the 33 clusters identified in a fetal sex-specific manner (**Table2, Fig.4E-F**).

**Table 2.**
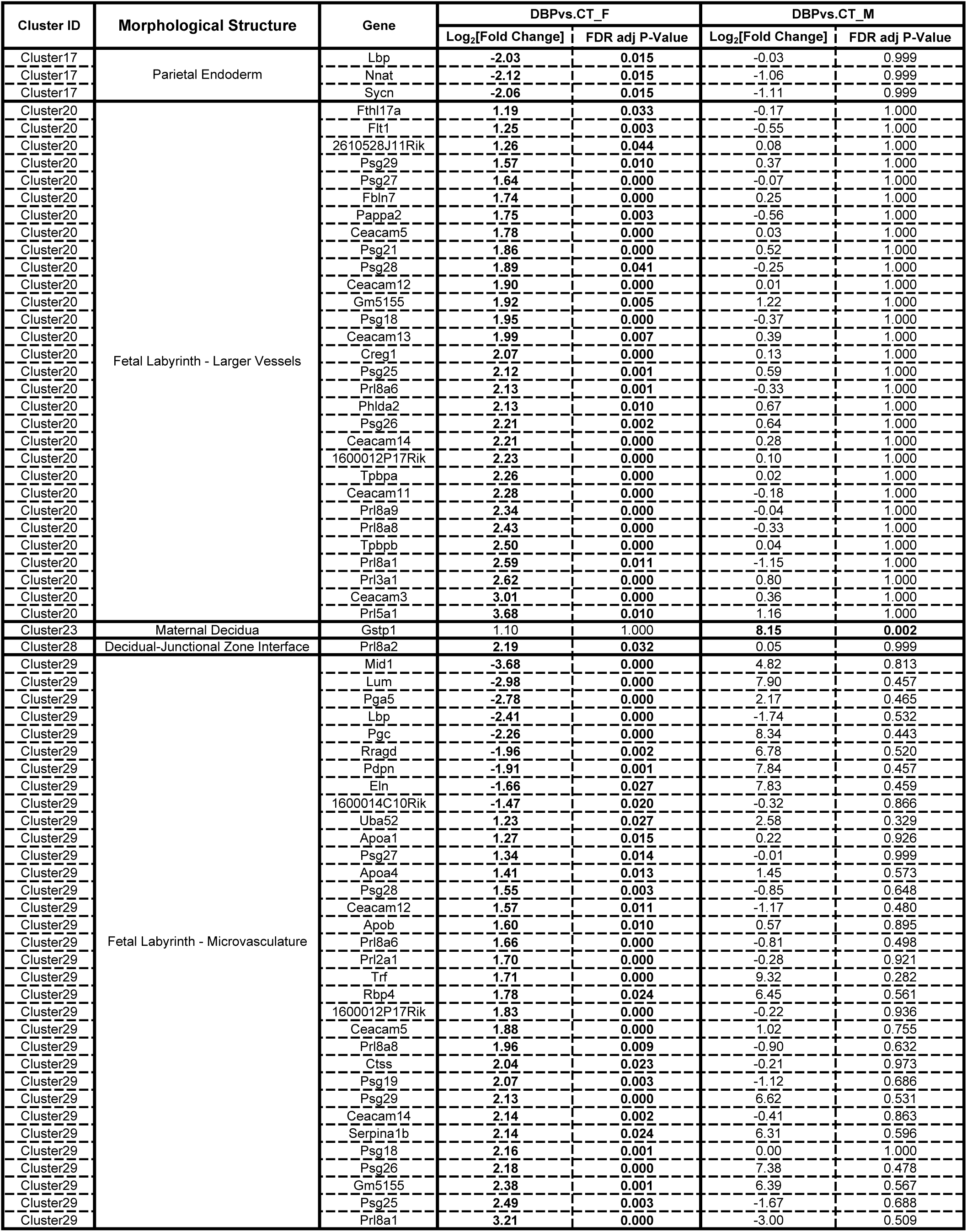
Prenatal DBP Exposure Dysregulated Genes in Spatially Resolved Placental Structures.

In male placentas, prenatal DBP exposure significantly up-regulated only one gene (Glutathione S-Transferase Pi 1 [*Gstp1*], an indicator of xenobiotic detoxification and cellular defense against oxidative stress activity^65, 66^) in the cluster 23 (Maternal Decidua) and did not affect the gene expression profiles of any other placental regions.

In female placentas, prenatal DBP exposure dysregulated significantly more genes than their male counterparts with the most significant effect seen in placental labyrinth vasculature areas. Specifically, in cluster 17 (Parietal Endoderm), the prenatal DBP exposure significantly down-regulated 3 genes in female placentas (Lipopolysaccharide-Binding Protein [*Lbp*]: linked to inflammatory response^67^; Neuronatin [*Nnat*]: associated with fetal development and metabolism^68^; Syncollin [*Sycn*], associated with lipid transport^69^). In cluster 20 (Fetal Lyrinth - Larger Vessels), prenatal DBP exposure up-regulated 30 genes in female placentas, which are enriched with genes associated with critical diseases and biological functions (such as chronic inflammatory disorder, cardiomyopathy, fibrosis of heart, senescence of cells, and protein kinase cascade; **Table.3** and **Table.S1**). In cluster 28 (Decidual-Junctional Zone Interface), prenatal DBP exposure up-regulated 1 gene (Prolactin Family 8, Subfamily A, Member 2 [*Prl8a2*], a member of the prolactin family that supports pregnancy recognition and maintenance^70^) in female placentas. In cluster 29 (Fetal Labyrinth - Microvasculature), the prenatal DBP exposure dysregulated 33 genes (24 up-regulated and 9 down-regulated) in female placenta that are enriched with genes associated with critical diseases and biological functions (such as cardiomyopathy, angiogenesis, fibrosis, inflammatory response, diabetes mellitus, fatty acid metabolism, apoptosis, development of body trunk, dystrophy of muscle, and metabolism of reactive oxygen species only; **Table.4** and **Table.S2**). The prenatal DBP exposure induced most significant gene expression changes in female placental labyrinth vascular areas (cluster 20 [larger vessels] and cluster 29 [microvasculature]). There are 14 genes that were commonly up-regulated in both the fetal labyrinth larger vessel and microvascular clusters, including carcinoembryonic antigen-related pregnancy-specific glycoproteins^71^ (*Ceacam5*, *Ceacam12*, *Ceacam14*, *Gm5155* [also known as *Ceacam23*], *Psg25*, *Psg26*, *Psg27*, *Psg28*, *Psg29*), prolactin-family genes^72^ (*Prl8a1*, *Prl8a6*, *Prl8a8*, *Psg18*), and 1600012P17Rik (a late gestation placenta-specific long noncoding RNA^73^).

**Table 3.**
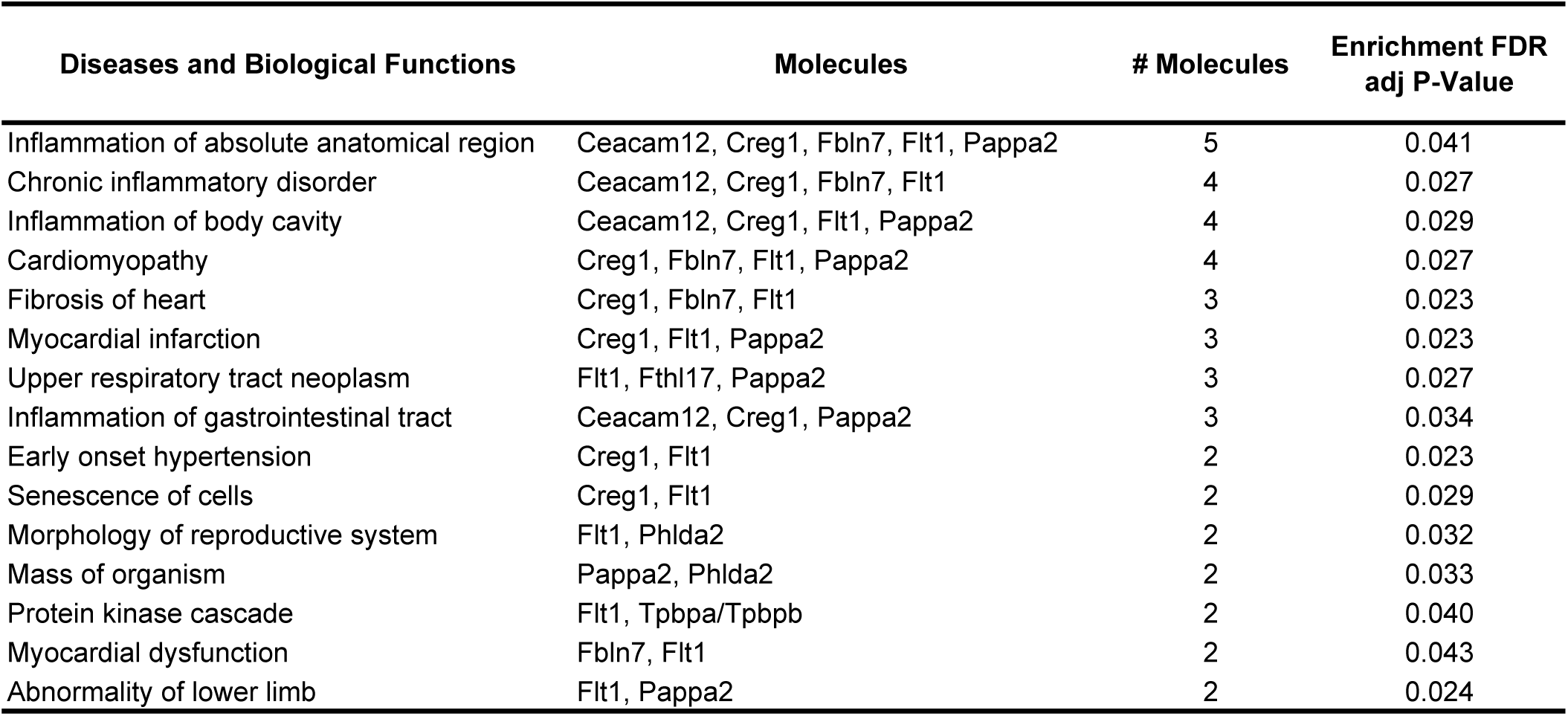
Top Enriched IPA Diseases and Biological Functions in Prenatal DBP Exposure Dysregulated Genes in Cluster 20 of Female Placentas [DBP vs. CT_F].

| **Diseases and Biological Functions** | **Molecules** | **# Molecules** | **Enrichment FDR**<br>**adj P-Value** |
| --- | --- | --- | --- |
| Inflammation of absolute anatomical region | Ceacam12, Creg1, Fbln7, Flt1, Pappa2 | 5 | 0.041 |
| Chronic inflammatory disorder | Ceacam12, Creg1, Fbln7, Flt1 | 4 | 0.027 |
| Inflammation of body cavity | Ceacam12, Creg1, Flt1, Pappa2 | 4 | 0.029 |
| Cardiomyopathy | Creg1, Fbln7, Flt1, Pappa2 | 4 | 0.027 |
| Fibrosis of heart | Creg1, Fbln7, Flt1 | 3 | 0.023 |
| Myocardial infarction | Creg1, Flt1, Pappa2 | 3 | 0.023 |
| Upper respiratory tract neoplasm | Flt1, Fthl17, Pappa2 | 3 | 0.027 |
| Inflammation of gastrointestinal tract | Ceacam12, Creg1, Pappa2 | 3 | 0.034 |
| Early onset hypertension | Creg1, Flt1 | 2 | 0.023 |
| Senescence of cells | Creg1, Flt1 | 2 | 0.029 |
| Morphology of reproductive system | Flt1, Phlda2 | 2 | 0.032 |
| Mass of organism | Pappa2, Phlda2 | 2 | 0.033 |
| Protein kinase cascade | Flt1, Tpbpa/Tpbpb | 2 | 0.040 |
| Myocardial dysfunction | Fbln7, Flt1 | 2 | 0.043 |
| Abnormality of lower limb | Flt1, Pappa2 | 2 | 0.024 |

**Table 4.**
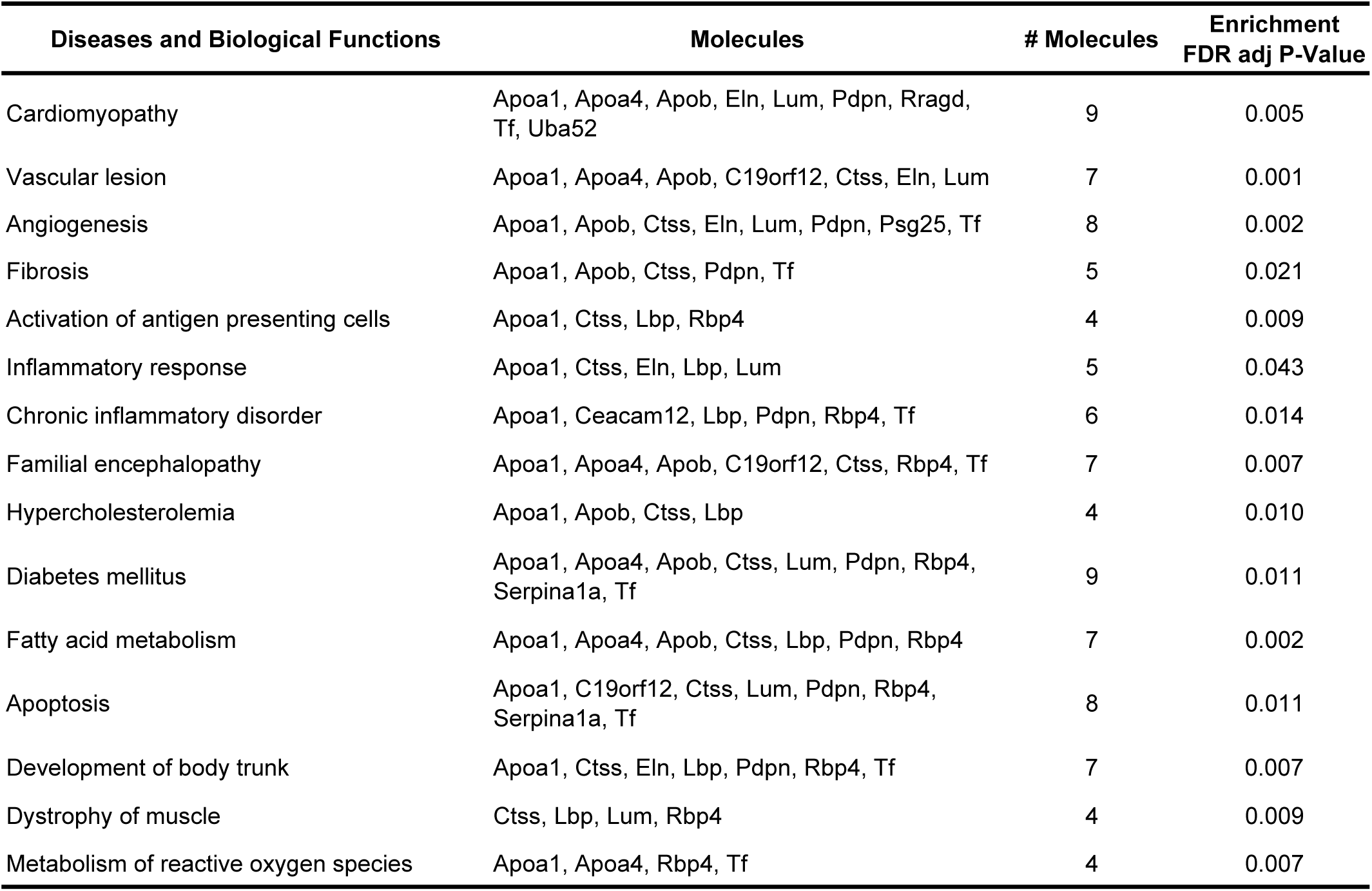
Top Enriched IPA Diseases and Biological Functions in Prenatal DBP Exposure Dysregulated Genes in Cluster 29 of Female Placentas [DBP vs. CT_F].

### Prenatal DBP Exposure Fetal Sex-Specifically Dysregulates Placental Lipid Profiles

Across the 12 classes of lipid molecules examined, only phosphatidylcholine (PC) was significantly elevated in male placentas from DBP groups compared with CT group, while no significant differences observed in female placentas (**Fig.5A**). In addition, the total abundance of PC was significantly higher in male than female placentas from DBP group, while there were no differences observed between male and female placenta from CT group. No class-level abundance differences were observed in other lipid molecules classes examined, including ceramides (Cer), dihydroceramides (dhCer), sphingoid bases and phosphates (Sphingoid_base), glucosylceramides (GlcCer), lactosylceramides (LacCer), diacylglycerols (DAG), phosphatidylethanolamines (PE), lysophosphatidylcholines (LPC), lysophosphatidylethanolamines (LPE), lysophosphatidylserines (LPS), acylcarnitines (AC).

**Figure. 5.**
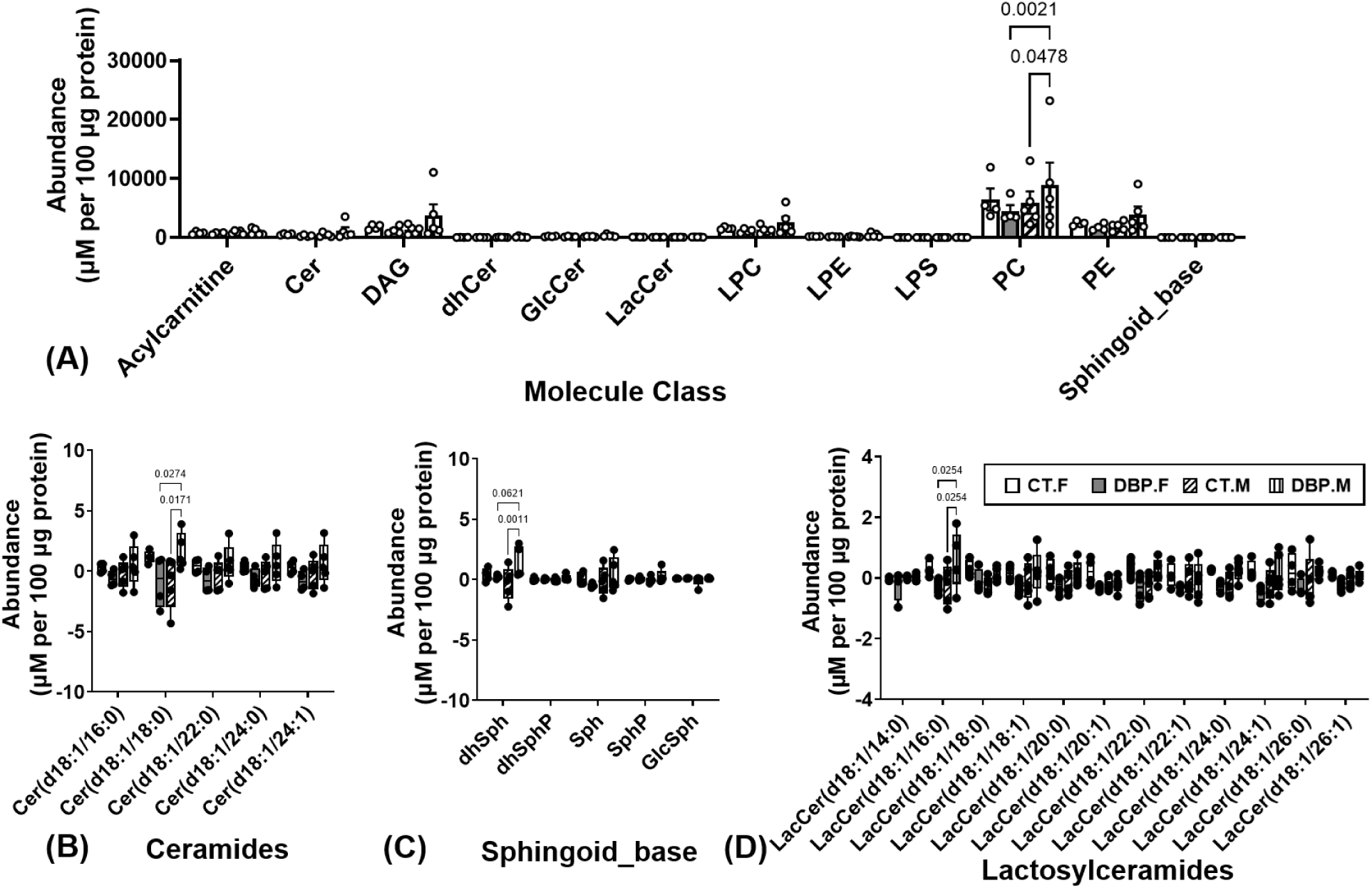
Prenatal low-dose DBP Exposure Fetal Sex-Specifically Dysregulates Placental Lipid Profiles of PC Class (A) as well as Ceramides(B), Sphingoid base (C), and Lactosylceramides (D) at Molecules. Differ between groups: P<0.05 (n=5 per fetal sex/group).

At the individual molecule level, prenatal DBP exposure elevated 3 molecules out of the 136 molecules assessed in a fetal sex-specifical manner. Specifically, prenatal DBP exposure elevated the abundance of C18 ceramide (Cer(d18:1/18:0)), dihydrosphingosines (dhSph), and C16 lactosylceramide (LacCer(d18:1/16:0)) in male placentas, but did not change in female placentas, compared to CT groups.

### Prenatal DBP Exposure Reduces GD18.5 PVECs Outgrowth Capacity

Primary PVECs preparations isolated from GD18.5 female and male control placentas (CT-F and CT-M) exhibited comparable in vitro outgrowth capacities, with cell growth observed in 40% (8 out of 20) of PVECs preparations in both CT-F and CT-M groups (**Fig.6A**). Prenatal DBP exposure reduced the proportion of PVECs preparations exhibiting growth in both sexes, with a more pronounced decrease seen in PVECs from male placentas. Specifically, cell growth was observed in 29.4% (5 out of 17) of DBP-F and 17.6% (3 out of 17) of DBP-M PVECs preparations, respectively, indicating potentially reduced in vitro proliferative capacity following prenatal DBP exposure.

**Figure. 6.**
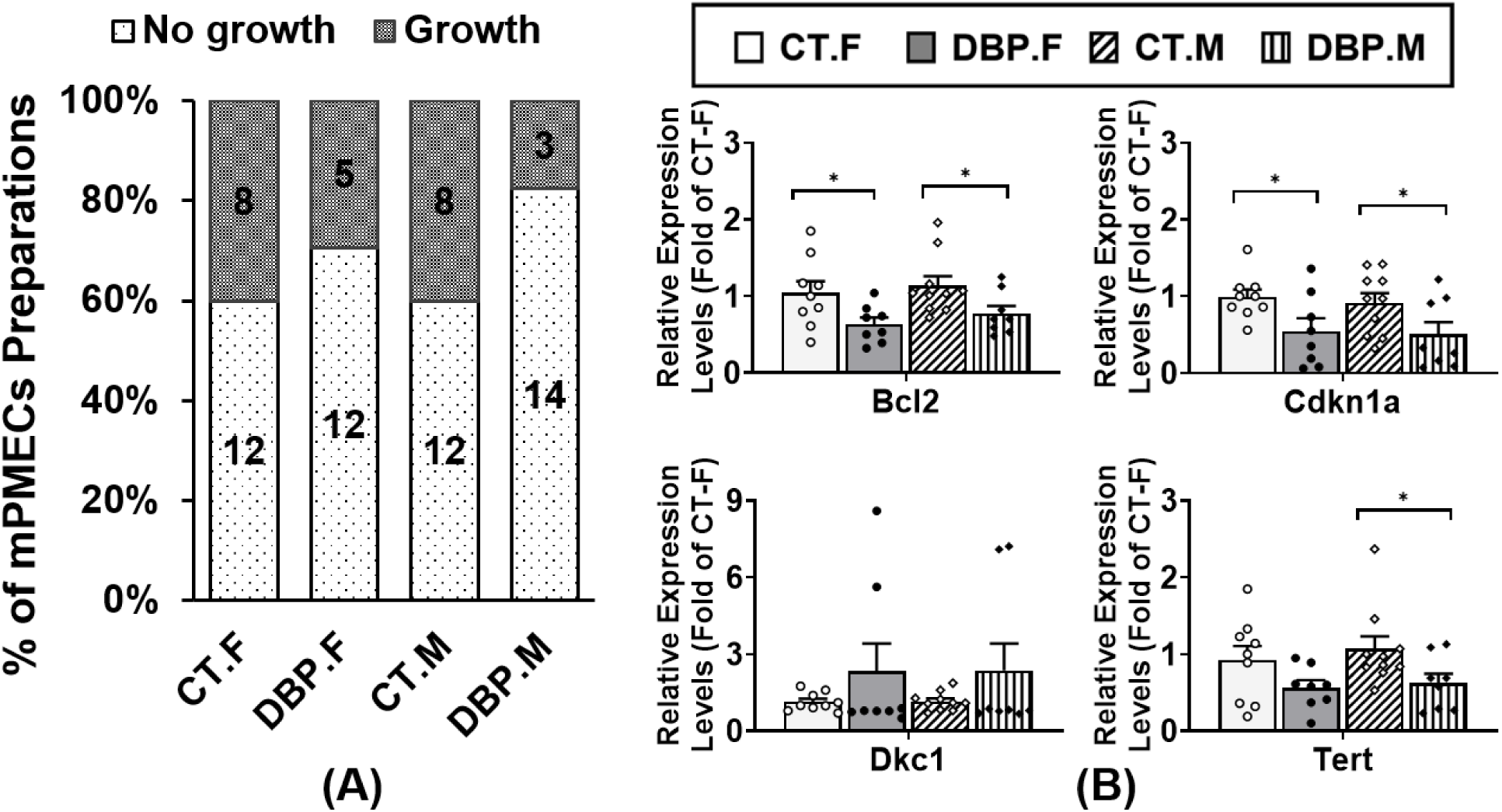
Prenatal low-dose DBP Exposure Fetal Sex-Specifically Dysregulates PVECs Outgrowth Capacities (A) and Whole Placental Expression of Anti-apoptotic, Senescence, and Telomere-Maintenance Related Genes (B). (A): CT-F: n=20 PVMCs from 8 dams, DBP-F: n=17 PVMCs from 7 dams, CT-M: n=20 PVMCs from 8 dams, DBP-M: n=17 PVMCs from 7 dams; (B): CT-F: n=9 dams, DBP-F: n=8 dams, CT-M: n=10 dams, DBP-M: n=8 dams. **\***Differ between groups (P<0.05).

### Prenatal DBP Exposure Dysregulates Anti-apoptotic, Senescence, and Telomere-Maintenance Related Genes in GD18.5 Placentas

The female and male placentas from control animals (CT-F and CT-M) exhibited comparable relative gene expression levels of *Bcl2* (anti-apoptosis marker^60^), *Cdkn1a* (senescence/cell-cycle arrest marker^61^), and *Dkc1* (telomerase stability marker^62^), and *Tert* (catalytic subunit of telomerase^63^). Prenatal DBP exposure significantly reduced the mRNA expression of *Bcl2* (decreased 36.5% in DBP-F and 22.5% in DBP-M) and *Cdkn1a* (decreased 44.2% in DBP-F and 41% in DBP-M) in GD18.5 placentas. Prenatal DBP exposure did not significantly change the *Dkc1* mRNA expression in both female and male placentas at GD18.5, although variability was observed in DBP-F and DBP-M placentas. Compared to control groups, *Tert* mRNA expression was significantly reduced by 44.9% in DBP-M placentas but was not significantly altered in DBP-F placentas at GD18.5.

## Discussion

In this study, we have demonstrated that prenatal exposure to an environmentally relevant low-dose of DBP (0.1µg/kg/day) disrupts fetal-placental vascular function during pregnancy in CD-1 mice. Although this DBP exposure did not significantly alter maternal uterine artery or vein hemodynamics, it dysregulated fetoplacental vascular indices during pregnancy and was associated with an increased proportion of low placental efficiency among male, but not female, fetuses at GD18.5. We further showed for the first time that low-dose prenatal DBP exposure induces sex-specific remodeling of the placental spatial transcriptomic profiles, particularly within labyrinth vasculature, and placental lipid abundance in a fetal sex-specific manner. In addition, we observed impaired outgrowth capacity of primary PVECs and fetal sex-specific dysregulation of anti-apoptotic, senescence, and telomere-maintenance related genes in GD18.5 placentas from fetuses with prenatal low-dose DBP exposure. Collectively, these data indicate that environmentally relevant low-dose prenatal DBP exposure compromise placental vascular function and endothelial proliferative potential in association with disrupted placental lipid metabolism. These findings provide new evidence that fetal-placental vasculature is a sensitive target of low-dose phthalate exposure.

The DBP exposure dose used in this study (0.1µg/kg/day) falls within the lower range of estimated human environmental exposure and is known to not induce systemic toxicity or altered weight gain pattern in both non-pregnancy and pregnant female CD-1 mice^27, 29, 30, 45, 46^. This low exposure dose was specifically chosen to evaluate the impact of environmentally relevant prenatal DBP exposure on fetal-placental development and function independent of the confounding obesogenic effects reported at higher DBP doses^10–12^.

Our high-frequency ultrasound data in maternal uterine arteries and veins showed that although 30 days of low-dose DBP exposure does not significantly affect the uterine vascular function in non-pregnant mice, it dysregulates maternal uterine vascular adaptation during mid gestation. As mid gestation is a critical period for placental development, this dysregulation in vascular adaptation could adversely affect placental development and function, which is supported by our high-frequency ultrasound data in fetal umbilical arteries. Consistent with previous report that DBP can cross the maternal-fetal interface and is associated with fetal growth restriction^16, 22^, our data showed that prenatal DBP exposure has more dramatic effects on the fetal vascular function than the maternal vasculature. We observed that prenatal DBP exposure reduced the umbilical artery RI at mid-gestation and attenuated the physiological EDV elevation from mid- to late-gestation, indicating dysregulated gestational adaptation of fetoplacental vascular tone. Together with previous report that DBP induces ex vivo vasorelaxation in human umbilical arteries^74^, these findings indicate that DBP exposure disrupts fetal-placental vasculature development and function during pregnancy. This finding is consistent with previous report in humans showing that prenatal DBP exposure (indicated by elevated urinary concentration of mono-n-butyl phthalate [MBP, primary metabolite of DBP] during late gestation) is associated with elevated blood pressure in children at 3-7 years old^75^.

Prenatal low-dose DBP exposure dysregulates the distributions of fetal weight, placental weight, and placental efficiency at GD18.5 in a fetal sex-specific manner. Prenatal DBP exposure shifted fetal weight, placental weight, and placental efficiency of female conceptuses towards intermediate category while shifted male conceptuses towards low fetal-placental efficiency category. These data indicating that male conceptuses are more susceptible to prenatal low-dose DBP exposure-associated impairment of placental efficiency. This finding is supported by human studies showing that prenatal DBP exposure (indicated by elevated urinary MBP levels) is associated with increased total placental surface area and placental thickness as well as less efficient fetal-placental vascular tree at delivery, with stronger association seen in male placentas^76, 77^.

Although previous studies in humans have linked prenatal DBP exposure (urinary MBP) to altered placental gene expression^38–42^, causality of these changes remains unclear due to multi-phthalate co-exposure and the heterogeneous cell compositions of placental tissues used in bulk transcriptomic analysis. Additionally, placental vascular function depends on precisely organized cellular architecture and cell-cell interactions which cannot be captured by bulk tissue transcriptomic profiling. In the present study, we performed spatial transcriptomic profiling analysis on female and male GD18.5 mice placentas from control and DBP exposed animals, to dissect prenatal DBP exposure-induced transcriptional remodeling within distinct placental compartments within the late-gestation placentas, thereby linking compartment-specific molecular responses to altered placental vascular development and function.

Our spatial transcriptomic profiling data revealed that prenatal low-dose DBP exposure dysregulates the placental transcriptomic profile in a morphological structure- and fetal sex-specific manner. *Gstp1* encodes glutathione S-transferase pi 1, a phase II detoxification enzyme involved in cellular defense against electrophilic and oxidative stress^65, 66^. In humans, elevated glutathione levels and glutathione peroxidase activity in decidual and chorionic villous placental tissues have been linked to hypertensive pregnancy complications (pre-eclampsia and hemolysis, elevated liver enzymes, low platelets [HELLP] syndrome)^78^. The male-specific upregulation of *Gstp1* observed in the maternal decidua (cluster 23) area in placentas from DBP exposed dams indicates a maternal decidual-localized activation of detoxification and cellular defense response against DBP exposure in male placentas^78^. Together with our observation that prenatal DBP exposure reduces placental efficiency exclusively in male fetuses, these data indicate an insufficient defense response in the male placentas in response to DBP exposure.

Prenatal DBP exposure dysregulated 68 genes among 5 clusters (cluster 17, 20, 23, 28, 29) in female placentas. The majority of these prenatal DBP exposure dysregulated genes (92.6%, 63 out of 68) in female placentas were localized to fetal labyrinth vascular regions, with 30 genes altered in the larger-vessels (cluster 20) and 33 genes altered in the microvascular (cluster 29). Together with our high-frequency ultrasound data, these findings provide strong evidence that prenatal low-dose DBP exposure directly affects placental vascular development and function.

The observed female-specific labyrinth vasculature-enriched transcriptional response indicates that the female fetal placental labyrinth vasculature is particularly sensitive to prenatal low-dose DBP exposure. Notably, 13 out of the 14 commonly up-regulated genes in fetal labyrinth larger vessels and microvasculature in female placentas are encoding carcinoembryonic antigen-related pregnancy-specific glycoproteins and prolactin-family genes that are associated with maternal-fetal immune modulation, angiogenic regulation, and pregnancy-related endocrine/paracrine adaptation^71, 72^. In addition, the differentially dysregulated genes in cluster 20 and cluster 29 both enriched with genes associated with inflammation and cardiomyopathy dysfunction (such as cardiomyopathy). This shared transcriptional signature suggests that prenatal DBP exposure induced a coordinated placental response across different fetal vascular compartments within the labyrinth. Prenatal DBP exposure also induces compartment-specific effects within the female fetal labyrinth vasculature, with senescence-related genes dysregulated only in larger vessels (Cluster 20), whereas lipid-metabolism related genes primarily dysregulated in microvascular (Cluster 29). This finding agrees with our previous report^27^ that the prenatal low-dose DBP exposure does not affect the lipid deposition in maternal decidual region in all placentas, while decreases lipid deposition in the fetal labyrinth regions in female, but not in male, placentas. These data indicating that these prenatal DBP exposure dysregulated lipid metabolism-associated genes identified in the female fetal labyrinth microvasculature region may contribute to the decreased lipid deposition observed in female placental fetal labyrinth. Together with our observation that prenatal low-dose DBP exposure adversely affects the fetal-placental efficiency in male conceptuses but shifts female conceptuses away from the extreme low- and high-efficiency categories, these female fetal labyrinth vascular region-specific changes reflect a compensational adaptation that preserves fetal-placental function.

Our lipid profiling analysis reveals that prenatal DBP exposure selectively disrupts placental lipid homeostasis in male, but not in female, placentas. Interestingly, this fetal sex-specific lipid profile dysregulation shows an inverse pattern to our spatial transcriptomic findings. Female placentas exhibit much more significant transcriptional remodeling in the placental fetal labyrinth vascular compartment while maintaining lipid homeostasis, whereas male placentas show minimal transcriptomic adaptation while exhibiting marked changes in lipid homeostasis. Together, these data suggest that male and female placentas employ different strategies in response to prenatal DBP exposure, with critical implications for sex-specific cardiovascular programming.

Phosphatidylcholine (PC) is the primary class of structural phospholipids in mammalian cell membranes and serve as precursors for bioactive lipid mediators implicated in endothelial dysfunction, vascular inflammation, and atherosclerosis^79, 80^. Elevated PC and dysregulated PC metabolism is linked to cardiovascular disease, metabolic syndrome, and insulin resistance^81, 82^. The accumulation of PC exclusively in male placentas without significant transcriptomic changes, indicates a failed transcriptional adaptation rather than an active compensatory response. This interpretation aligns with the established paradigm that male placentas prioritize growth maintenance rather than protective adaptations when exposed to environmental stressors^83, 84^.

Accumulation of ceramide (Cer) and lactosylceramide (LacCer) are reported to induce vascular endothelial dysfunction by reducing nitric oxide (NO) bioavailability and increasing oxidative stress^85–88^. The male placenta-specific accumulation of C18 ceramide [Cer(d18:1/18:0)] and C16 lactosylceramide [LacCer(d18:1/16:0)] in prenatal DBP exposed animals reflects a lipotoxic profile associated with increased oxidative stress and endothelial dysfunction. This pattern resembles placental oxidative stress and lipid profile dysregulation reported in pregnancy complications (preeclampsia and fetal growth restriction) characterized by placental dysfunction and increased offspring cardiovascular disease risk^89–93^. Together with report in humans showing that prenatal DBP exposure is associated with elevated triglycerides and increased risks of non-alcoholic fatty liver disease during young adulthood in offspring^94^, the lipid homeostasis dysregulation observed in this study represents a novel mechanism linking prenatal DBP exposure to developmental programming of offspring cardiometabolic disease.

The reduced outgrowth capacities observed in PVECs from prenatal DBP exposed animals provides direct evidence that low-dose DBP exposure impairs the proliferative potential of placental vascular endothelial cells. Compared to the baseline outgrowth ratio among PVECs from CT-F (40%) and CT-M (40%) placentas, reduced outgrowth potential was observed in both DBP-F (29.4%) and DBP-M (17.6%) PVECs. This finding aligns with the decreased *Bcl2* and *Cdkn1a* expression in both DBP-F and DBP-M placentas compared to their control counterparts. *Bcl2* is an anti-apoptosis marker that promotes cell survival^60^, while *Cdkn1a* is a marker for stress-induced senescence and DNA damage repair^61^. This observation is consistent with previous report in CD-1 mice that DBP exposure decreases *Bcl2*, increases *Cdkn1a* expression in antral follicles, and induces cell cycle arrest and atresia^37^. The reduction of both anti-apoptotic and stress-adaptive cell cycle control related genes could contribute to the reduced outgrowth potential under in vitro culture seen in PVECs from DBP exposed animals possibly through inducing cell cycle arrest or senescense.

The more pronounced PVECs outgrowth reduction in DBP-M compared to DBP-F groups is consistent with the Cer(d18:1/18:0) and LacCer(d18:1/16:0) accumulation observed exclusively in DBP-M placentas, as Cer and LacCer species are reported to induce endothelial cell growth arrest and apoptosis^95, 96^. This finding is further supported by the male-specific reduction (44.9% decrease) of *Tert* expression in DBP exposed placentas which indicative of impaired telomere maintenance and accelerated telomere shortening^63, 97^. In DBP-M placentas, the absence of spatially specific transcriptomic changes despite significant whole-placenta *Tert* down-regulation indicates a diffused, global expression reduction across multiple cell types instead of a confined, specific placental compartment. This finding indicates a widespread failure of transcriptional adaptation which is consistent with the ceramide accumulation seen and impaired endothelial proliferation potential in DBP-M placentas. Given the essential role of *Tert* in telomere maintenance in endothelial cells^97^, and previous report linking reduced *Tert* activity to cellular senescence and endothelial dysfunction^98^, this male placenta-specific *Tert* reduction indicating that the prenatal low-dose DBP exposure may contribute to the impaired placental endothelial function potential through telomere-maintenance disruption. The preservation of *Tert* expression in DBP-F placentas, may represent a sex-specific protective mechanism that partially preserved female placentas against ceramide accumulation and endothelial dysfunction. Given the reported role of TERT in maintaining mitochondrial function and protecting against oxidative and ischemic injury^99^, the enrichment of myocardial dysfunction-related genes in female placental labyrinth vascular regions may reflect activation of shared mitochondrial/vascular stress-response pathways. Together, these findings suggest that female placentas undergo transcriptional remodeling in response to prenatal DBP exposure that may protect against lipotoxicity and endothelial dysfunction.

In conclusion, our data have demonstrated that prenatal low-dose DBP exposure dysregulates placental vascular function and lipid homeostasis in a fetal sex-specific manner. Female placentas exhibited fetal labyrinth vasculature-specific transcriptomic adaptations and relatively preserved placental efficiency and endothelial function. In contrast, male placentas exhibited minimum transcriptomic adaptation, together with compromised placental efficiency and endothelial function associated with lipotoxic lipid profile.

## Acknowledgments

We would like to thank Dr. Xiaosong Liu for technical assistance.

---

**Table S1.** Enriched Diseases and Biological Functions in Prenatal DBP Exposure Dysregulated Genes in Cluster 20 of Female Placentas [DBP vs. CT_F].

| Diseases and Biological Functions | Molecules | # Molecules | Enrichment FDR<br>adj P-Value |
| --- | --- | --- | --- |
| Inflammation of absolute anatomical region | Ceacam12, Creg1, Fbln7, Flt1, Pappa2 | 5 | 0.041 |
| Chronic inflammatory disorder | Ceacam12, Creg1, Fbln7, Flt1 | 4 | 0.027 |
| Cardiomyopathy | Creg1, Fbln7, Flt1, Pappa2 | 4 | 0.027 |
| Inflammation of body cavity | Ceacam12, Creg1, Flt1, Pappa2 | 4 | 0.029 |
| Fibrosis of heart | Creg1, Fbln7, Flt1 | 3 | 0.023 |
| Myocardial infarction | Creg1, Flt1, Pappa2 | 3 | 0.023 |
| Upper respiratory tract neoplasm | Flt1, Fthl17, Pappa2 | 3 | 0.027 |
| Inflammation of gastrointestinal tract | Ceacam12, Creg1, Pappa2 | 3 | 0.034 |
| Benign pelvic disease | Creg1, Flt1, Phlda2 | 3 | 0.038 |
| Senescence of cells | Creg1, Flt1 | 2 | 0.029 |
| Morphology of reproductive system | Flt1, Phlda2 | 2 | 0.032 |
| Mass of organism | Pappa2, Phlda2 | 2 | 0.033 |
| Protein kinase cascade | Flt1, Tpbpa/Tpbpb | 2 | 0.040 |
| Myocardial dysfunction | Fbln7, Flt1 | 2 | 0.043 |
| Abnormality of lower limb | Flt1, Pappa2 | 2 | 0.024 |
| Enlargement of heart | Creg1, Fbln7, Flt1 | 3 | 0.023 |
| Early onset hypertension | Creg1, Flt1 | 2 | 0.023 |
| Interstitial fibrosis | Fbln7, Flt1 | 2 | 0.023 |
| G1/S phase transition | Creg1, Flt1 | 2 | 0.023 |
| Neovascularization | Creg1, Flt1 | 2 | 0.024 |
| Autophosphorylation of protein | Fbln7, Flt1 | 2 | 0.024 |
| Failure of kidney | Fbln7, Flt1 | 2 | 0.024 |
| Hypertrophy of heart | Creg1, Flt1 | 2 | 0.031 |
| Inflammation of liver | Creg1, Flt1 | 2 | 0.040 |
| Chronic kidney disease | Fbln7, Flt1 | 2 | 0.042 |
| Dysfunction of heart | Creg1, Flt1 | 2 | 0.044 |
| Size of lesion | Creg1, Flt1 | 2 | 0.049 |
| Ear-nose-throat cancer | Flt1, Fthl17, Pappa2 | 3 | 0.023 |
| Gastroenteritis | Ceacam12, Creg1, Pappa2 | 3 | 0.031 |
| Smooth muscle tumor | Creg1, Flt1, Phlda2 | 3 | 0.023 |
| Anaplastic carcinoma | Flt1, Fthl17 | 2 | 0.047 |
| Calcinosis | Fbln7, Flt1 | 2 | 0.027 |
| Carcinoid tumor | Flt1, Pappa2 | 2 | 0.038 |
| Cell proliferation of sarcoma cell lines | Creg1, Flt1 | 2 | 0.024 |
| Invasive gastric cancer | Flt1, Fthl17 | 2 | 0.023 |
| Laryngeal squamous cell carcinoma | Flt1, Pappa2 | 2 | 0.046 |
| Liver cholangiocarcinoma | Flt1, Pappa2 | 2 | 0.023 |
| Liver Damage | Creg1, Flt1 | 2 | 0.024 |
| Pancreatic endocrine tumor | Flt1, Pappa2 | 2 | 0.032 |
| Pharyngeal squamous cell carcinoma | Flt1, Pappa2 | 2 | 0.023 |
| Uterine leiomyoma | Creg1, Phlda2 | 2 | 0.032 |

**Table S2.**
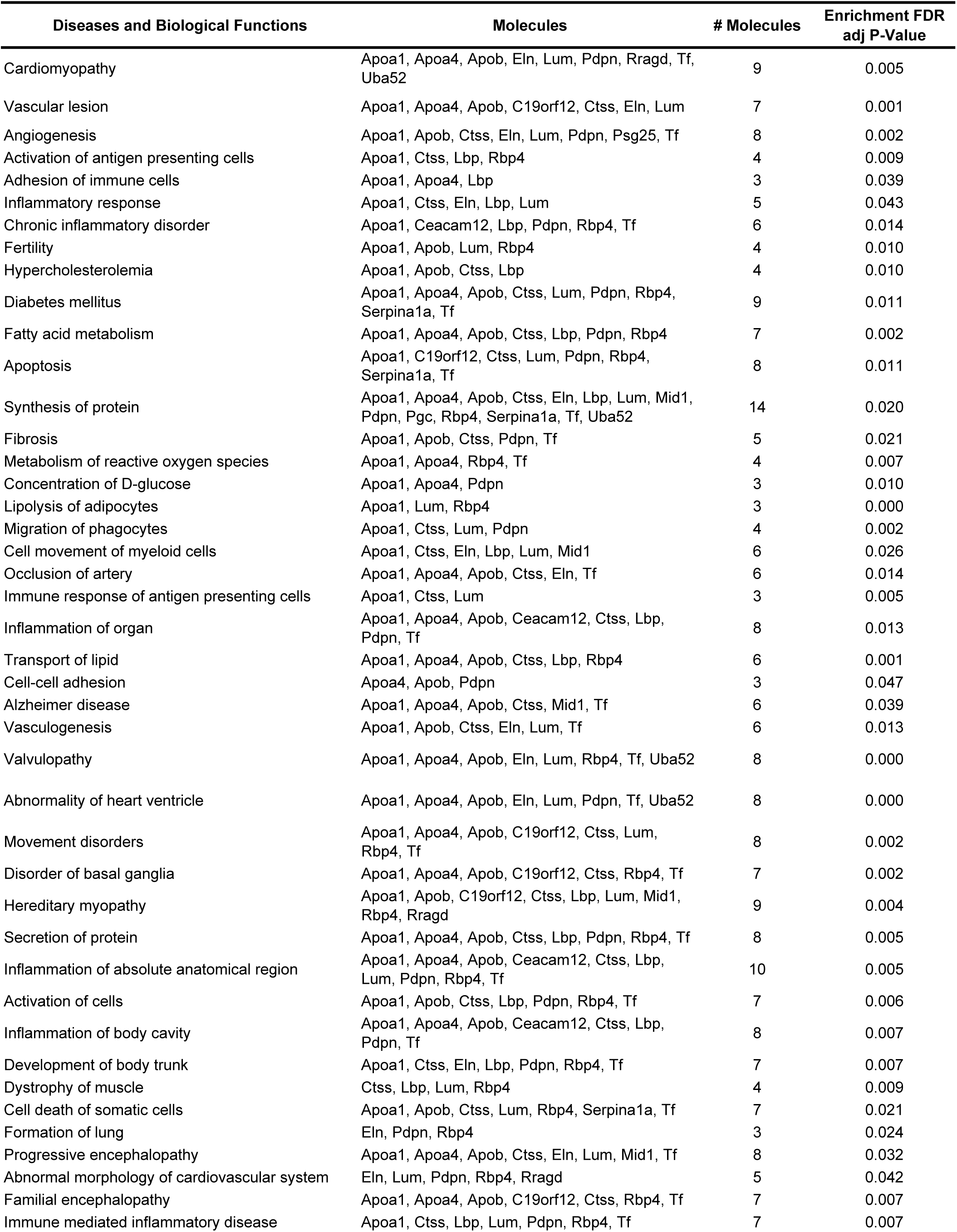
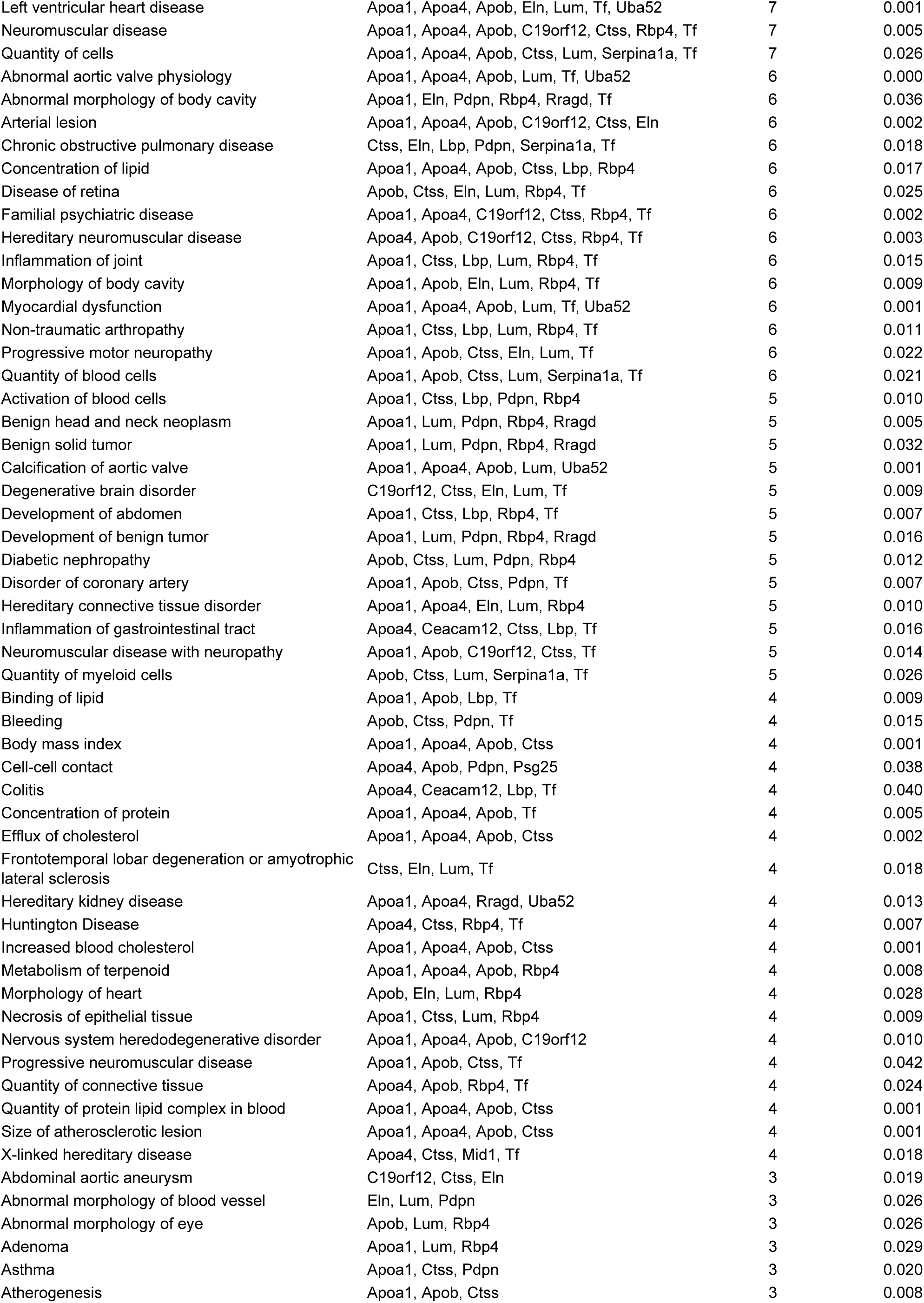
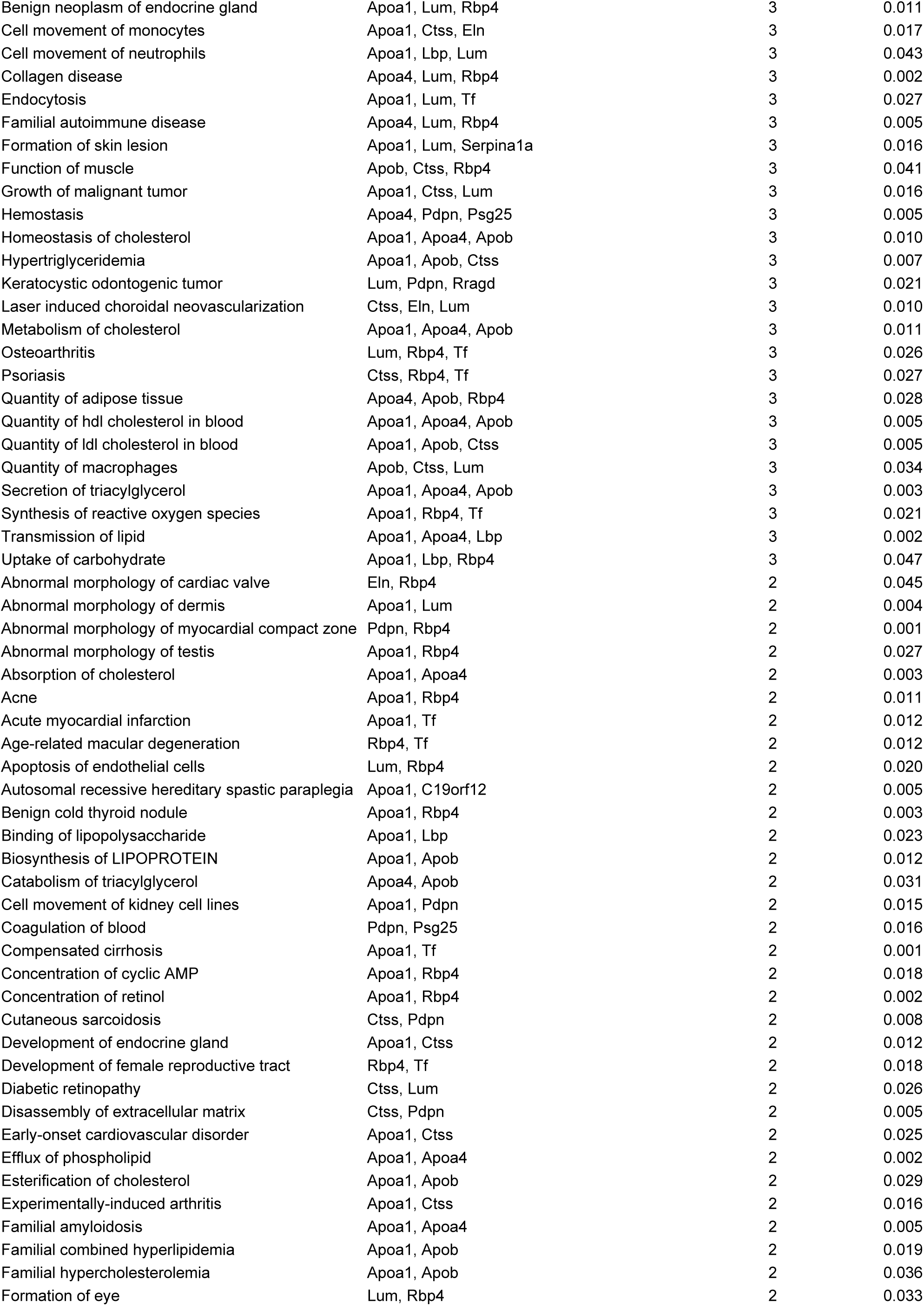
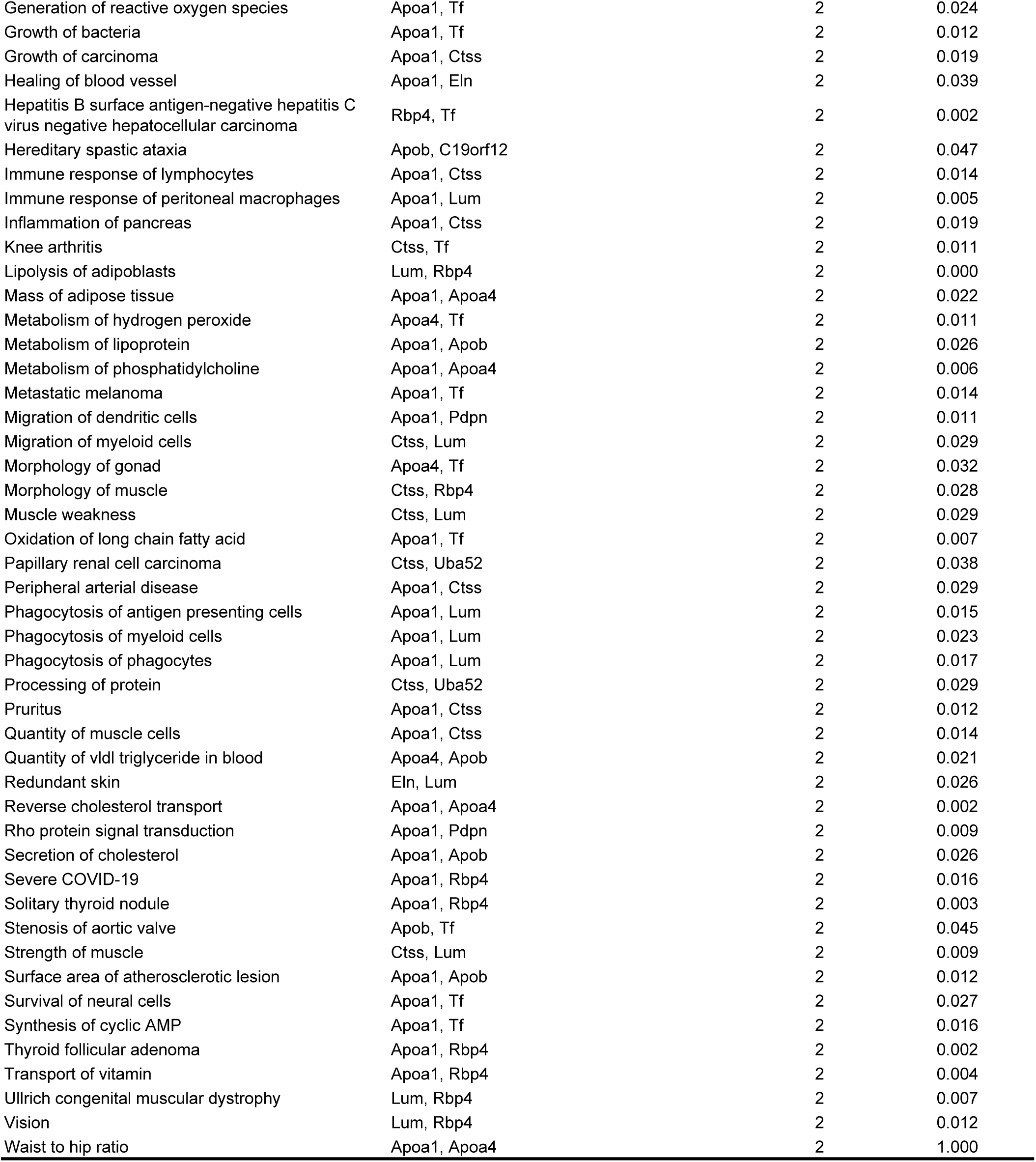
Enriched IPA Diseases and Biological Functions in Prenatal DBP Exposure Dysregulated Genes in Cluster 29 of Female Placentas [DBP vs. CT_F].

